# Donor-specific assemblies enhance somatic structural variant detection in complex genomic regions

**DOI:** 10.64898/2026.02.20.707061

**Authors:** Taralynn Mack, Jiadong Lin, Luyao Ren, Min-Hwan Sohn, Anna Minkina, Youngjun Kwon, DongAhn Yoo, Yang Sui, Katherine M Munson, Kendra Hoekzema, F Kumara Mastrorosa, Melanie Sorensen, Marcelo Ayllon, Kaitlyn A Sun, Nidhi Koundinya, Jeffrey Ou, Michelle D Noyes, Adriana Sedeño-Cortés, Amy Leonardson, Caitlin N Jacques, Chris Oliviera, Christian D Frazar, Christina Zakarian, Dana M Jensen, Elliott G Swanson, Erica Ryke, J Thomas Kolar, Jane Ranchalis, Lila Sutherlin, Mitchell R Vollger, Kelsey Loy, Meranda M Pham, Meng-Fan Huang, Natalie YT Au, Patrick M Nielsen, Sean R McGee, Shane Neph, Stephanie Bohaczuk, Tristan Shaffer, Vea Freeman, Yizi Mao, Benjamin Cohen Stillman, Matthew Richardson, Joshua D Smith, Jeffrey M Weiss, Nancy L Parmalee, Chia-Lin Wei, James T Bennett, Andrew B Stergachis, Evan E Eichler

## Abstract

Structural variants (SVs) contribute substantially to genomic variation and disease, but detecting somatic SVs (sSVs) remains difficult due to reference bias, mosaicism, and enrichment in repetitive regions. Linear reference genomes, like GRCh38 and CHM13, do not fully capture individual genomic structure, which can obscure true somatic variation. Donor-specific assemblies (DSAs) generated from the same genome where sSVs are being assayed provide a personalized alternative, yet their performance for sSV detection has not been systematically assessed. As part of the Somatic Mosaicism across Human Tissues (SMaHT) Network, we benchmark a DSA for sSV discovery in the COLO829 melanoma cell line with a matched normal sample from the same individual. We compare sSV detection across GRCh38, CHM13, and COLO829BL_DSA using three sSV callers (DELLY, Severus, and Sniffles2) and multiple long-read sequencing platforms and validate candidate calls through visual inspection of aligned reads. COLO829BL_DSA identifies 1.8-fold more validated sSVs than linear references, in regions both shared with GRCh38 and CHM13 and unique to COLO829BL_DSA. Variants detected only with COLO829BL_DSA are often found in satellite and other repeat-rich regions that are difficult to resolve using standard references, including two variants in cancer-associated genes not identified by either linear reference. We further identify caller-, platform-, and region-specific filtering strategies that substantially improve precision, offering a more scalable path toward high-confidence sSV detection. Overall, these results underscore the utility of DSAs, alongside practical strategies for their analysis, in improving sSV detection.

## INTRODUCTION

Structural variants (SVs), defined as genetic alterations of at least 50 base pairs in length, have a well-established role in cancer development (Cosenza et al. 2022; Zhang et al. 2019; Yang and Yang 2023; Lopez et al. 2020). Beyond cancer, SVs have recently been implicated in a broad range of additional diseases, including neurodegenerative disorders (Li et al. 2025; Miller et al. 2021) and age-related clonal hematopoiesis (Loh et al. 2018). However, detecting SVs remains technically challenging. Germline SVs are often large, nonrandomly distributed, and enriched in repetitive and structurally complex genomic regions that are difficult to resolve using short-read sequencing, which shows reduced sensitivity and a higher false discovery rate for SV detection in these regions (Sedlazeck et al. 2018). The landscape of somatic structural variants (sSVs) is less well understood. Detecting sSVs is further complicated by their mosaic nature and potential low variant allele frequency, along with challenges specific to somatic variant calling, such as tumor purity, tumor heterogeneity, and alignment ambiguity in repetitive regions (Gong et al. 2020).

Traditional linear human reference genomes (e.g., GRCh38) represent consensus sequences from multiple donors (International Human Genome Sequencing Consortium 2001). They often fail to capture individual-specific SVs, especially within repetitive or highly polymorphic regions—precisely the regions of the genome that are most mutable (Porubsky et al. 2025). This reference bias can obscure true variation and limit the sensitivity of sSV detection. More recently, nearly complete telomere-to-telomere (T2T) human reference genomes (e.g., CHM13) have been produced from a single source material (Nurk et al. 2022). Yet, germline variants present in an individual may still be entirely missed when they occur in genomic regions absent from that reference (Chaisson et al. 2015b).

The development of long-read sequencing technologies has significantly improved the resolution of SVs, enabling more comprehensive detection across all regions of the genome (Chaisson et al. 2015a). In parallel, donor-specific assemblies (DSAs)—personalized reference genomes built by *de novo* assembly of an individual donor’s own sequencing data rather than by alignment to a pre-existing reference—have emerged to enhance SV detection and improve mapping in repetitive genomic regions, allowing both somatic and germline variants to be readily distinguished (Nurk et al. 2022; Sohn et al. 2026). DSAs are typically built by *de novo* assembly of long-read sequencing data, often combined with Hi-C data for haplotype phasing, to produce a phased, diploid, near-T2T genome. Because this genome reflects the donor’s own sequence directly, including structural variation within segmental duplications and satellite repeats largely absent from standard linear references such as GRCh38 or CHM13, it reduces reference bias and improves the sensitivity and specificity of somatic variant calling (Somatic Mosaicism across Human Tissues Network 2026). Recent work has demonstrated the utility of using a DSA to detect somatic single-nucleotide variation (Porubsky et al. 2025; Sohn et al. 2026), and, more recently, sSVs (Zhang et al. 2025), highlighting the potential of this approach while underscoring the need for broader benchmarking.

Here, as part of the benchmarking efforts of the Somatic Mosaicism across Human Tissues (SMaHT) Network, we assess the utility of using a DSA for detecting sSVs in the well-characterized COLO829 melanoma cell line, which includes matched tumor (COLO829) and normal (COLO829BL) samples. We generated sequencing data from three complementary long-read sequencing platforms: PacBio high-fidelity (HiFi), standard Oxford Nanopore Technologies (STD-ONT), and ultra-long ONT (UL-ONT). These datasets were analyzed with three sSV callers, DELLY (Rausch et al. 2012), Severus (Keskus et al. 2026), and Sniffles2 (Smolka et al. 2024), which were chosen due to their performance in previous long-read cancer genome benchmarking studies (Aydin et al. 2025). We use these callers to identify five classes of sSVs: insertions (INS), deletions (DEL), duplications (DUP), inversions (INV), and breakends (BND). Insertions add new genetic sequence, deletions remove sequence, and duplications create additional copies of existing sequence. Inversions flip the orientation of a sequence segment, while breakends represent points of discontinuity in the genome that can indicate translocations or other more complex rearrangements (Feuk et al. 2006). The aims of this study were to (i) benchmark sSV detection using a DSA against the GRCh38 and CHM13 references across three long-read sequencing platforms and three sSV callers, (ii) identify sSVs that are accessible only with a DSA, and (iii) define caller-, platform-, and region-specific filtering strategies that improve the precision of DSA-based sSV callsets.

## RESULTS

### Construction of a high-quality donor-specific assembly capturing near-T2T completeness

In addition to using the standard genome references for sSV detection, we constructed a DSA from HiFi, UL-ONT, and Hi-C data produced from the normal lymphoblastoid source material (COLO829BL) and the details of this are described elsewhere (Sohn et al. 2026). Briefly, we applied the assembly algorithm Verkko 2.1 (Antipov et al. 2025) to generate a phased diploid genome where two haplotypes were assembled near T2T. The resulting assembly harbored less than 50 gaps per haplotype with an estimated single-nucleotide variant error rate of <1 in a million base pairs, with at least nine non-acrocentric chromosomes represented T2T in at least one haplotype. We observed that, except for Chromosome 21, all acrocentric regions were assembled continuously from the p-arm to the q-arm in both haplotypes with the exception of the rDNA cluster. We also investigated regions of potential errors based on depth of realigned reads and estimated 23.9 Mbp of potential collapse and 6.9 Mbp of misassembly. These numbers are comparable to near-T2T genome assemblies released by international consortia, including the HPRC (Liao et al. 2023) and HGSVC (Logsdon et al. 2025). Comparison to GRCh38 identified 342.8 Mbp of sequence not present in the standard reference genome of which 61% of it was composed of satellite repeats and 22% represented segmental duplications unique to COLO829BL_DSA. We refer to this reference genome as COLO829BL_DSA to distinguish it from later assemblies of the tumor DNA which is used to validate more complex sSVs.

### sSV detection across references reveals higher sensitivity in COLO829BL_DSA

Three sSV callers (DELLY, Severus, and Sniffles2) were used to call sSVs using the COLO829 data aligned to three references (GRCh38, CHM13, and COLO829BL_DSA) using three sequencing data types PacBio HiFi, STD-ONT, and UL-ONT (**Figure 1A, Supplemental Table S1**). Across all sequencing data types and callers there were 13,975 total sSVs called for GRCh38, 24,396 for CHM13, and 18,061 for COLO829BL_DSA that were 50 bp or longer. Post-calling filtering was performed to remove both sequencing artifacts and germline events. Filtering sSVs in regions of known assembly artifacts or misassembly/collapse removed 7% of sSVs from GRCh38, 31% of CHM13 sSVs, and 14% of COLO829BL_DSA sSVs. Running Phased Assembly Variant (PAV) caller (Ebert et al. 2021) to detect germline SV differences between COLO829BL_DSA and the respective assemblies resulted in the removal of an additional 16% of GRCh38 variants and 10% of CHM13 sSVs as germline artifacts (**Methods**, **Supplemental Fig. S1A**).

**Figure 1.**
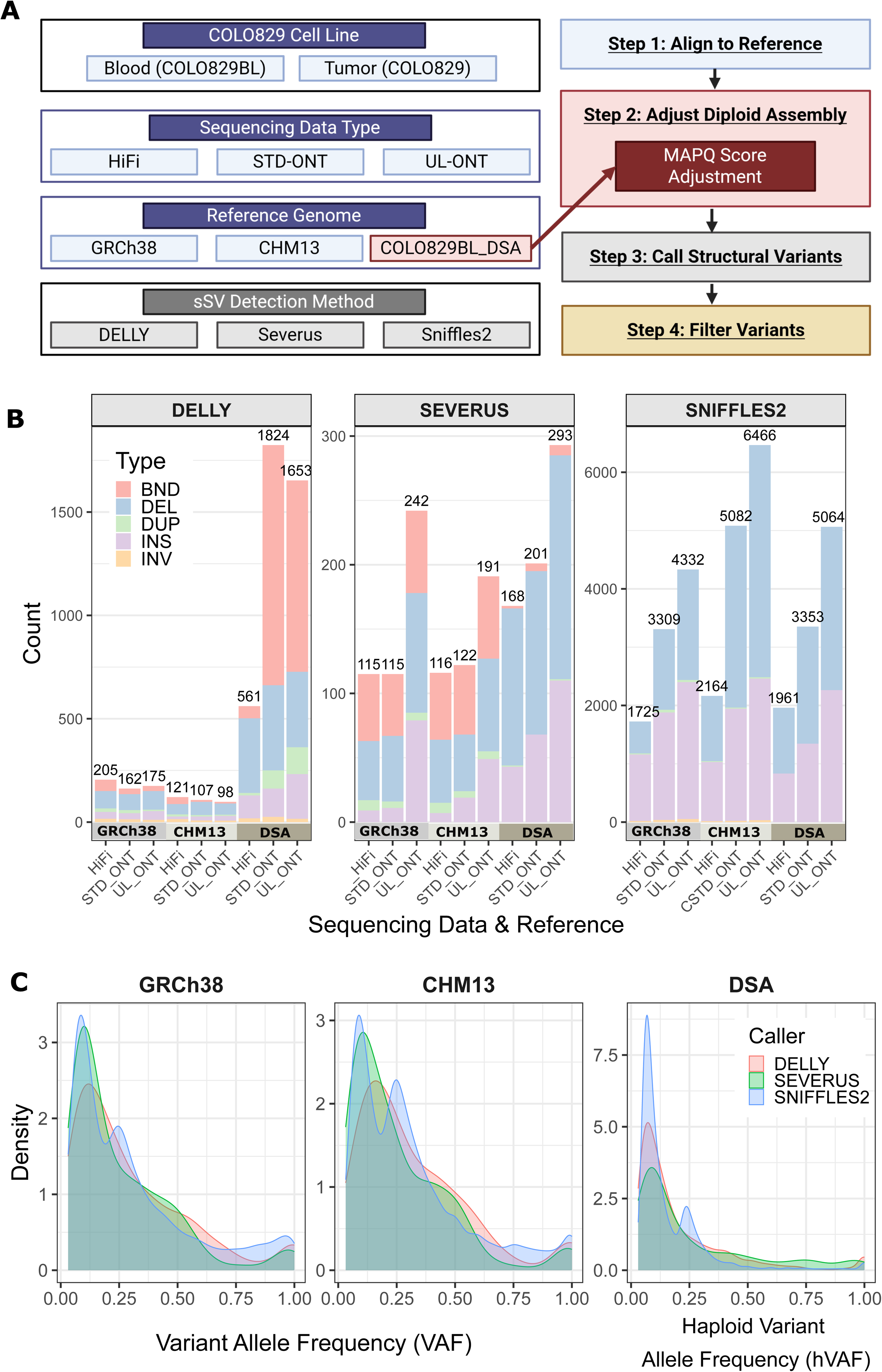
Somatic structural variant (sSV) discovery. **A)** Schematic depicting study design. PacBio high-fidelity (HiFi), standard Oxford Nanopore Technologies (STD-ONT), and ultra-long ONT (UL-ONT) sequencing data for COLO829BL and COLO829 were aligned to three reference genomes: GRCh38, CHM13, and COLO829BL_DSA. MAPQ values were manually adjusted for the COLO829BL_DSA calls to avoid variant dropout. Three sSV callers, DELLY (Rausch et al. 2012), Severus (Keskus et al. 2026), and Sniffles2 (Smolka et al. 2024) were used to detect somatic structural variation. **B)** Post-filtering sSV call counts for each caller, reference genome, and type of sequencing data. Color represents the type of sSV. Note sSV counts vary by more than an order of magnitude depending on the caller. **C)** Variant allele frequency (VAF) distributions for GRCh38 and CHM13, and haploid variant allele frequency (hVAF) for COLO829BL_DSA, shown for raw sSVs stratified by reference genome and caller. DSA, donor-specific assembly.

After the filtering steps, the remaining calls differed by both caller and reference: DELLY (GRCh38: 542, CHM13: 326, COLO829BL_DSA: 4,038), Severus (GRCh38: 472, CHM13: 429, COLO829BL_DSA: 662), and Sniffles2 (GRCh38: 9,366, CHM13: 13,712, COLO829BL_DSA: 10,378) (**Figure 1B, Supplemental Fig. S1B**). DELLY and Severus detected the most variants using COLO829BL_DSA, with up to a 15-fold increase in DELLY calls. Sniffles2 detected the most variants in CHM13. More variants were detected using ONT across all callers, especially UL-ONT.

The COLO829 dataset originates from a single tumor cell, meaning our results contain two classes of sSVs: those that are fixed in the tumor cell line (high variant allele frequency (VAF)) and those that arose during subsequent cell-culture expansion (low VAF). The parameters for all three sSV callers were set to detect both types, ensuring that we capture the full set of differences between COLO829BL and COLO829 (**Figure 1C**). COLO829BL_DSA calls are reported with haploid variant allele frequency (hVAF) since the sSVs are called from a diploid genome, and COLO829 ploidy has been reported (Sohn et al. 2026).

### Cross-caller and cross-platform sSV integration reveals COLO829BL_DSA-unique events

To produce a consensus sSV callset, we applied a two-stage integration strategy comprising (i) redundancy reduction and (ii) cross-caller and cross-sequencing-platform validation (**Figure 2A**). Using Truvari (English et al. 2022), the filtered variants were collapsed into one callset per reference genome regardless of sequencing data type (**Methods, Supplemental Fig. S1C**). Most variants were detected by only one platform, with 95% of calls unique to either HiFi or ONT (STD-ONT and/or UL-ONT) across all references, with ONT-only accounting for most variants (79%). ONT-only variants comprised 36% for DELLY, 63% for Severus, and 78% for Sniffles2, whereas HiFi-only variants represented a smaller fraction (36%, 13%, and 17%, respectively) (**Figure 2B**). COLO829BL_DSA continued to show the greatest sSV count for DELLY and Severus, while CHM13 had the greatest sSV count for Sniffles2 (**Figure 2B**).

**Figure 2.**
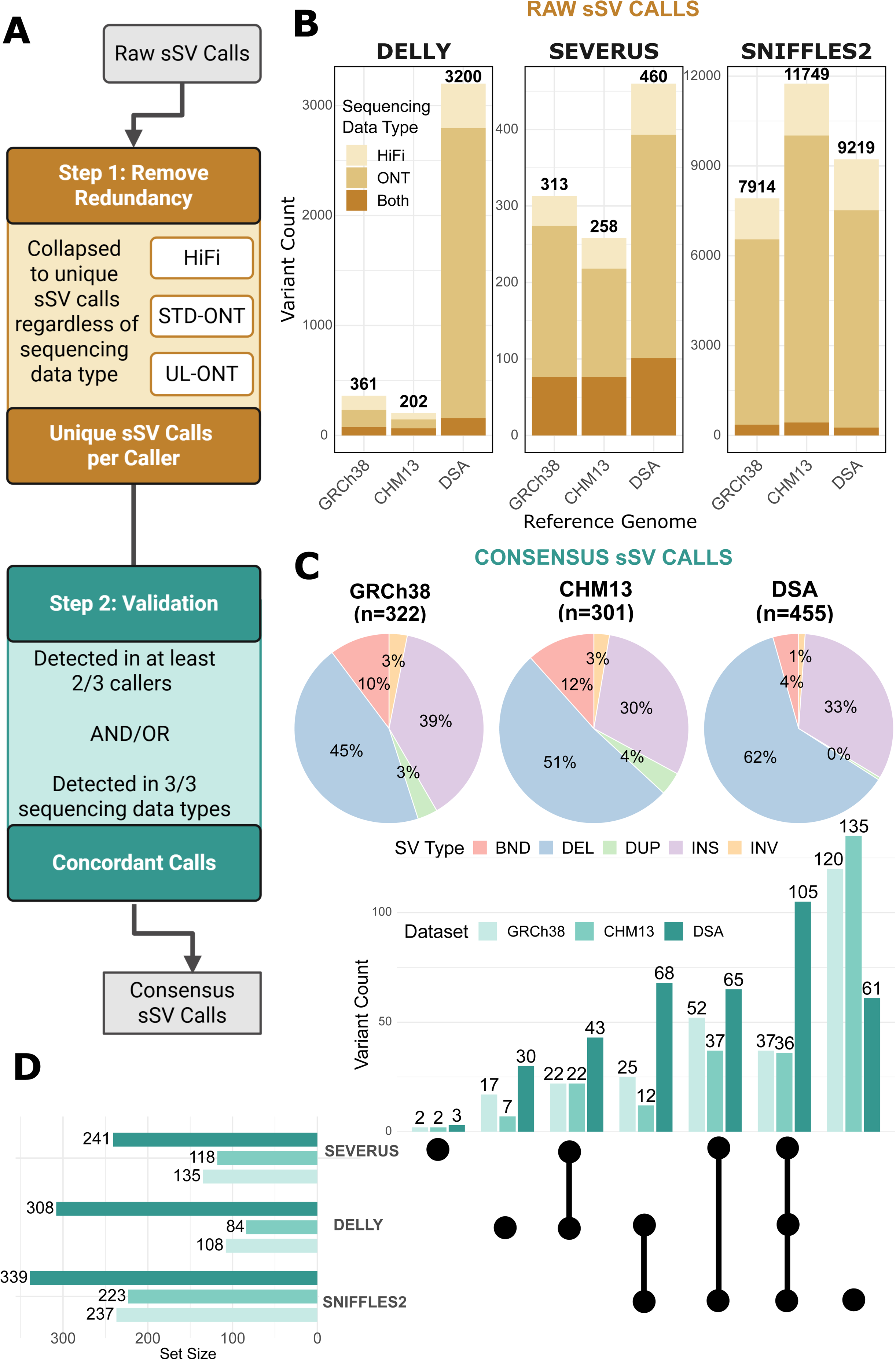
sSV callset integration. **A)** Schematic illustrating the process of integrating raw sSV calls, which involves removing redundancy by sequencing data type and validating each variant through detection by multiple callers or sequencing data types. **B)** Plot summarizing the number of unique variants per caller and reference genome after integrating all sequencing data types (HiFi, STD-ONT, UL-ONT). Bars are colored by the sequencing data types that detected each variant: HiFi, ONT (STD-ONT or UL-ONT), or both. **C)** Pie charts summarizing the number of variants in each reference genome detected by at least two of three sSV callers or by all three sequencing data types. Slice color indicates structural variant type: pink - BND (breakend), blue - DEL (deletion), green - DUP (duplication), purple - INS (insertion), and orange - INV (inversion). **D)** Upset plot indicating which callers detected each variant in the consensus callset and the total number of variants contributed by each sSV caller. Bar color corresponds to the reference genome.

After reduction, calls were integrated into one callset per reference genome. An sSV was included in the consensus dataset if it was detected by at least two of the three sSV callers, or if it was detected across all three sequencing data types for a single caller. This integration yielded 322 variants for GRCh38, 301 variants for CHM13, and 455 variants for COLO829BL_DSA. Insertions, deletions, breakends, duplications, and inversions were observed across all three references (**Figure 2C**). The largest category of variants in the consensus dataset (34%) was supported by only Sniffles2, which were later discovered to be all false positives, while the smallest category (<1%) consisted of variants detected exclusively by Severus. When considering COLO829BL_DSA calls alone, the largest fraction of variants was supported by all three sSV callers, as a smaller proportion of them are supported by Sniffles2 alone (**Figure 2D**). Within the consensus callset, the majority of variants were supported by all three sequencing platforms independently (70% GRCh38, 77% CHM13, and 52% DSA).

### Visual validation confirms that COLO829BL_DSA improves detection of true sSVs

Following the curation of the consensus dataset, we visually validated every non-BND variant by inspection of the underlying genome alignment and sequencing reads. There were 54 true positive variants for GRCh38, 51 for CHM13, and 95 for COLO829BL_DSA, corresponding to 20%, 20%, and 25% of the candidate variants, respectively. Validation rates differed substantially by genomic context: in non-TR regions, validation rates were 44% (GRCh38), 51% (CHM13), and 56% (DSA), whereas in TR regions they dropped sharply to 2%, 1%, and 16%, respectively (**Figure 3A**), reflecting the far greater difficulty of confidently validating calls in repetitive sequence regardless of reference assembly.

**Figure 3.**
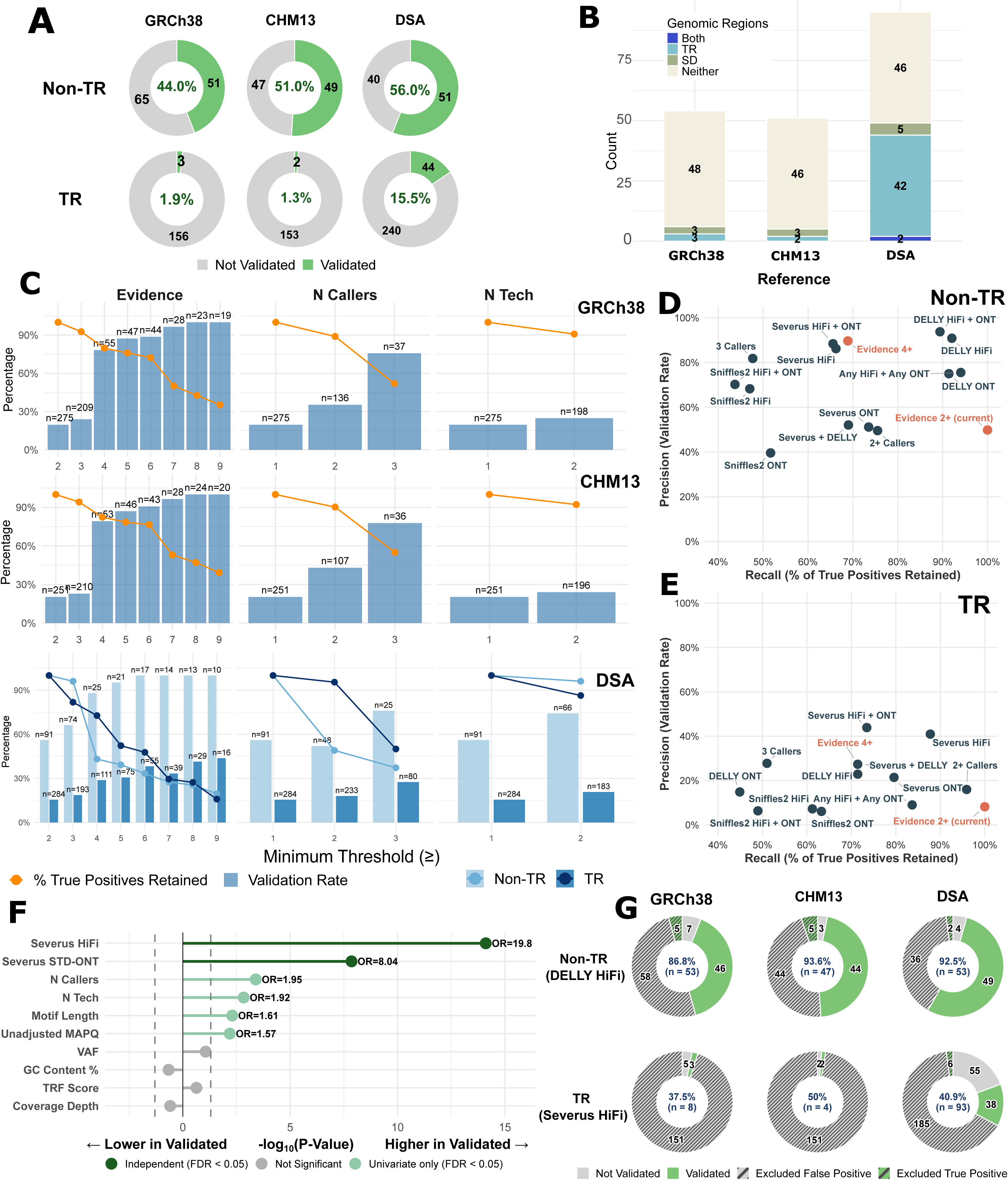
Reference comparisons and filtering improvements. **A)** Donut plots showing the proportion of validated (green) versus non-validated (gray) sSV calls from the consensus set within tandem repeat (TR)-associated and non-TR regions for each reference assembly (GRCh38, CHM13, DSA). Center labels indicate the overall validation rate, and segment labels show raw counts. **B)** Plot showing the number of sSVs mapping to tandem repeat (TR) or segmental duplication (SD) regions of the genome across the three different reference genomes. **C)** Validation rate (bars) and percentage of all true positives retained (lines) that would result from applying increasingly strict evidence, caller-count, or technology-count thresholds, shown separately for each assembly. Evidence score represents how many caller/sequencing data combinations showed support for that sSV. For DSA, results are further stratified by non-TR (light blue) and TR (dark blue) regions. Bar labels indicate the number of calls that would be retained at each threshold. **D,E)** Precision (validation rate) versus recall (percentage of true positives retained) under hypothetical filtering strategies based on individual callers, sequencing technologies, and combined evidence criteria, evaluated separately for non-TR (D) and TR (E) calls pooled across all three reference datasets. Points highlighted in orange indicate the current evidence-based filtering strategy (evidence ≥2) alongside an alternative threshold (evidence ≥4; see panel C), shown for comparison against caller- and technology-specific criteria to identify which strategy would offer the greatest improvement. **F)** Univariate association of candidate predictors with validation status among TR calls in the DSA dataset, identifying which additional features would be most informative for improving TR-specific filtering. Bars represent −log10(p-value), signed by direction of effect (right = higher in validated calls, left = lower); dashed lines indicate the significance threshold (p = 0.05). Odds ratios (ORs) are shown for predictors reaching statistical significance. Point/bar color indicates whether each predictor would provide independent value in a multivariate model (dark green), showed association only in univariate testing (light green), or showed no association (gray). **G)** Hypothetical filtering outcomes if non-TR calls were required to have DELLY HiFi support and TR calls were required to have Severus HiFi support, shown by reference dataset. Colored segments (green/gray) represent calls that would be retained under this criterion, split by validation status, while excluded segments (hashed) represent calls that would be filtered out, split by whether they were true or false positives (green/gray). Center labels indicate the validation rate (precision) that would be achieved among retained calls as well as the raw counts, illustrating the improvement over current practice.

Most true positive variants (87%) were called by both HiFi and ONT platforms (**Supplemental Fig. S2A**). As expected, variants supported by all three sequencing platforms validated at a higher rate than the consensus set overall in every reference (25% vs. 20% GRCh38, 23% vs. 20% CHM13, 37% vs. 25% DSA). Among individual platforms, HiFi-supported calls had the highest validation rate across all three references (25% GRCh38, 24% CHM13, 33% DSA), followed by STD-ONT and then UL-ONT. This gap widened substantially among platform-exclusive calls, with variants detected only by UL-ONT validating at just 3% (GRCh38), 3% (CHM13), and 4% (DSA), the lowest of any platform in every reference. The largest category of the visually validated variants (49%) was confirmed by all three sSV callers, while no visually validated variants were detected by Sniffles2 alone (**Supplemental Fig. S2B**).

Among validated variants, the vast majority of GRCh38 (89%) and CHM13 (90%) variants fell outside of tandem repeat (TR) and/or segmental duplication regions. In contrast, 46% of DSA-validated variants mapped within TR regions, consistent with the DSA’s improved ability to resolve variation in complex regions of the genome (**Figure 3B**). We also compared the MAPQ-adjusted and an unadjusted DSA callset: of the 95 validated true positives, 59 (62%) were only recovered after adjustment, at the cost of more false positives (precision dropped from 40% unadjusted to 25% adjusted).

To provide further validation, we also constructed a tumor-specific assembly (COLO829_DSA) with the existing long reads and compared it to COLO829BL_DSA (**Methods**). Although highly fragmented and incomplete, the resulting cancer cell line genome assembly had a contig N50 of 134.5 Mbp, with a QV of 57.9 based on HiFi reads. We compared COLO829_DSA versus COLO829BL_DSA and called sSVs using PAV (Ebert et al. 2021). Of the visually validated variants, 42% were supported by this assembly-to-assembly comparison, further evidence of the sSVs detected by the read-based callers. Of the variants missed by PAV, 43% occurred in regions absent from the tumor assembly, while the remaining 57% were largely explained by low hVAF (51%), inversions/duplications (11%), or satellite repeat context (26%), reflecting the intrinsic challenges of somatic variant detection. This is expected as PAV was designed to call germline variants and not somatic mutations; its use here, however, confirms high-frequency events at the assembly level.

Given that the overall validation rates were low across all three references, especially within TR regions, we next tested whether our current consensus criteria could be improved to yield a more precise callset for future work. We tested how precision and recall changed as a function of alternative evidence-score, caller-count, and technology-count thresholds (**Figure 3C**). Recall reflects the percentage of true positives within the consensus callset retained at each threshold, not all possible true positives across the raw callset. Across all three references, increasing the minimum evidence threshold improved precision at the cost of recall, with our current criterion (evidence ≥2) retaining the most consensus-set true positives but at comparatively low precision. An evidence threshold of ≥4, corresponding to multi-caller support, offered a reasonable balance for non-TR calls, improving precision substantially with only a modest recall trade-off. This did not hold for TR calls, however, which needed much higher evidence thresholds to reach comparable precision, and even then, at a steep cost to recall.

We next evaluated precision and recall for individual callers, technologies, and their combinations, separately for non-TR (**Figure 3D**) and TR (**Figure 3E**) sSV calls. DELLY HiFi retained the greatest number of true positives at strong precision in non-TR regions, while Severus HiFi did the same in TR regions. DELLY and Sniffles2 performed poorly on TR calls regardless of technology. Univariate and multivariate testing of candidate predictors of TR-call validation (**Figure 3F**) identified Severus HiFi (OR=19.8) and Severus STD-ONT (OR=8.04) support as the strongest independent predictors; other candidate features were associated with validation only in univariate testing or not at all.

Based on these results, we tested an alternative filtering strategy requiring DELLY HiFi support for non-TR calls and Severus HiFi support for TR calls (**Figure 3G**). This improved non-TR precision to 87% (GRCh38, n=53), 94% (CHM13, n=47), and 93% (DSA, n=53), and TR precision to 38% (GRCh38, n=8), 50% (CHM13, n=4), and 41% (DSA, n=93), though with reduced recall.

### COLO829BL_DSA uncovers unique sSVs missed by standard references

To compare sSV calls across the unique coordinate systems of the references, we implemented a graph-based approach (**Methods**) to remap variants. When all 95 visually validated sSVs from COLO829BL_DSA were converted to CHM13, 36 variants failed to convert using this approach. All 36 are unique to COLO829BL_DSA and would not be detected by the other reference genomes (**Figure 4A, Supplemental Table S2**). As expected, the majority of COLO829BL_DSA-unique variants mapped to repetitive regions, with 86% located in satellite repeats, and of those satellites, 75% corresponded to alpha satellites, including sSVs mapping to higher-order alpha satellite DNA (**Figure 4B-C**). sSVs were distributed across the chromosomes, with the highest number of sSVs detected on Chromosomes 7 and X (**Figure 4D**).

**Figure 4.**
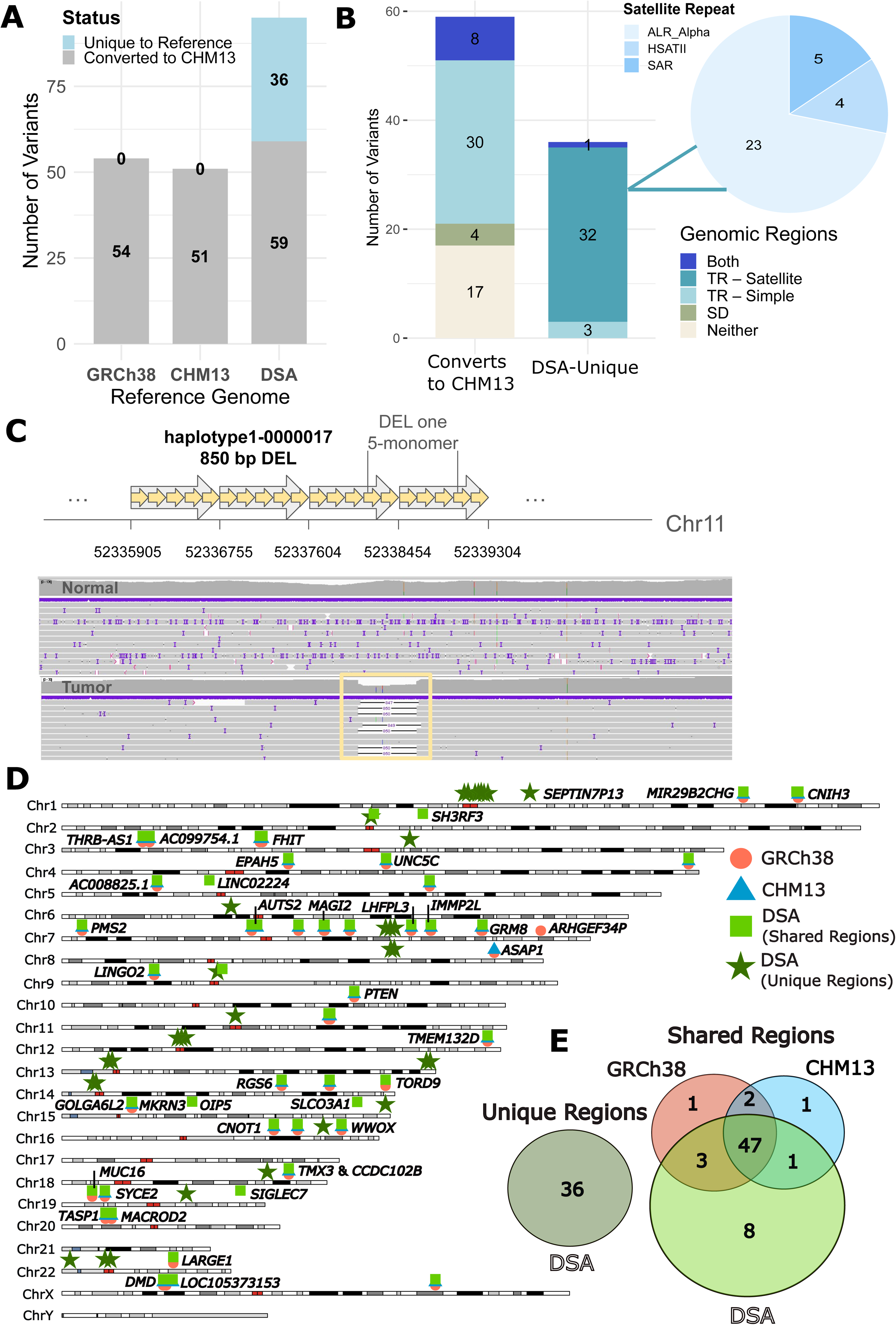
Sequence properties and genome-wide distribution of sSVs. **A)** Plot showing the number of sSV calls for each reference genome converted to CHM13 coordinates (displayed in Figure 3C,D) compared to the percentage that are unique to that reference genome (not displayed in Figure 3C,D). **B)** Genomic regions of COLO829BL_DSA variants for either COLO829BL_DSA variants converted to CHM13 or those unique to COLO829BL_DSA. For those unique to COLO829BL_DSA, a pie chart of the satellite subtypes is shown, with the vast majority as alpha satellites. **C)** Example of a deletion in a COLO829BL_DSA-unique alpha satellite region. **D)** Ideogram showing the location of sSV calls across the genome for all visually validated sSVs able to be mapped to CHM13 coordinates. Each reference is represented by a different color and shape, and overlap occurs when the shapes intersect. Because COLO829BL_DSA-unique SVs (green stars) lack direct CHM13 alignments, their positions are approximate, estimated from the nearest aligned flanking sequence. For any variants that overlap genes, the gene name is notated. **E)** Venn diagram displaying the number of overlapping sSV calls between GRCh38, CHM13, and COLO829BL_DSA when all calls are mapped to CHM13 coordinates (Shared Regions, n = 63), and variants unique to COLO829BL_DSA that occur in regions of the genome not represented in GRCh38 or CHM13 (Unique Regions, n = 36).

Among the 63 sSVs with true, shared coordinates across all three references (**Figure 4E**), 13% were detected using only COLO829BL_DSA, while 75% were detected by all three references. Four variants (6%) were not detected by COLO829BL_DSA and were instead detected by GRCh38 and/or CHM13. For these 63 sSVs, there was no significant difference in length or VAF between those detected only by COLO829BL_DSA and those shared across references.

We compared our results to Espejo Valle-Inclan et al. 2022, which reported 68 experimentally validated sSVs in COLO829 using a reference-based approach with PCR confirmation. Because their method relied on standard linear references, all 68 of their reported variants fall within genomic regions also accessible to GRCh38 (Shared Regions, **Figure 4E**). Restricting our comparison to the shared, mappable regions, our consensus callset recovered 85% of these variants when restricted to non-BND variants ≥50 bp.

To identify variants of potential functional consequence, we focused on sSVs mapping near or potentially disrupting protein-coding genes. Among the sSVs detected by all three references, 39 out of 47 are located within genes (**Table 1, Methods**), and of those, six impact the coding regions of the genes, *LHFPL3, PTEN, GOLGA6L2, CNOT1, TMX3-CCDC102B,* and *DMD.* Four of those, *LHFPL3* (Zhang et al. 2020), *PTEN* (Yehia et al. 2025; Khasarah et al. 2025)*, GOLGA6L2* (Sarker et al. 2025), and *TMX3-CCDC102B* (Luo et al. 2023; Si et al. 2022) have known links to cancer (**Table 1**).

Of those sSVs detected only by COLO829BL_DSA, 5/8 overlap genes and are intronic. Three of these overlap potential regulatory regions, *OIP5 (***Figure 5A**)*, SLCO3A1* (**Figure 5B**), and *SIGLEC7* (**Figure 5C**); *SIGLEC7* (Benthami et al. 2025; Thangaraj et al. 2025) has known links to cancer (**Table 2**).

**Figure 5.**
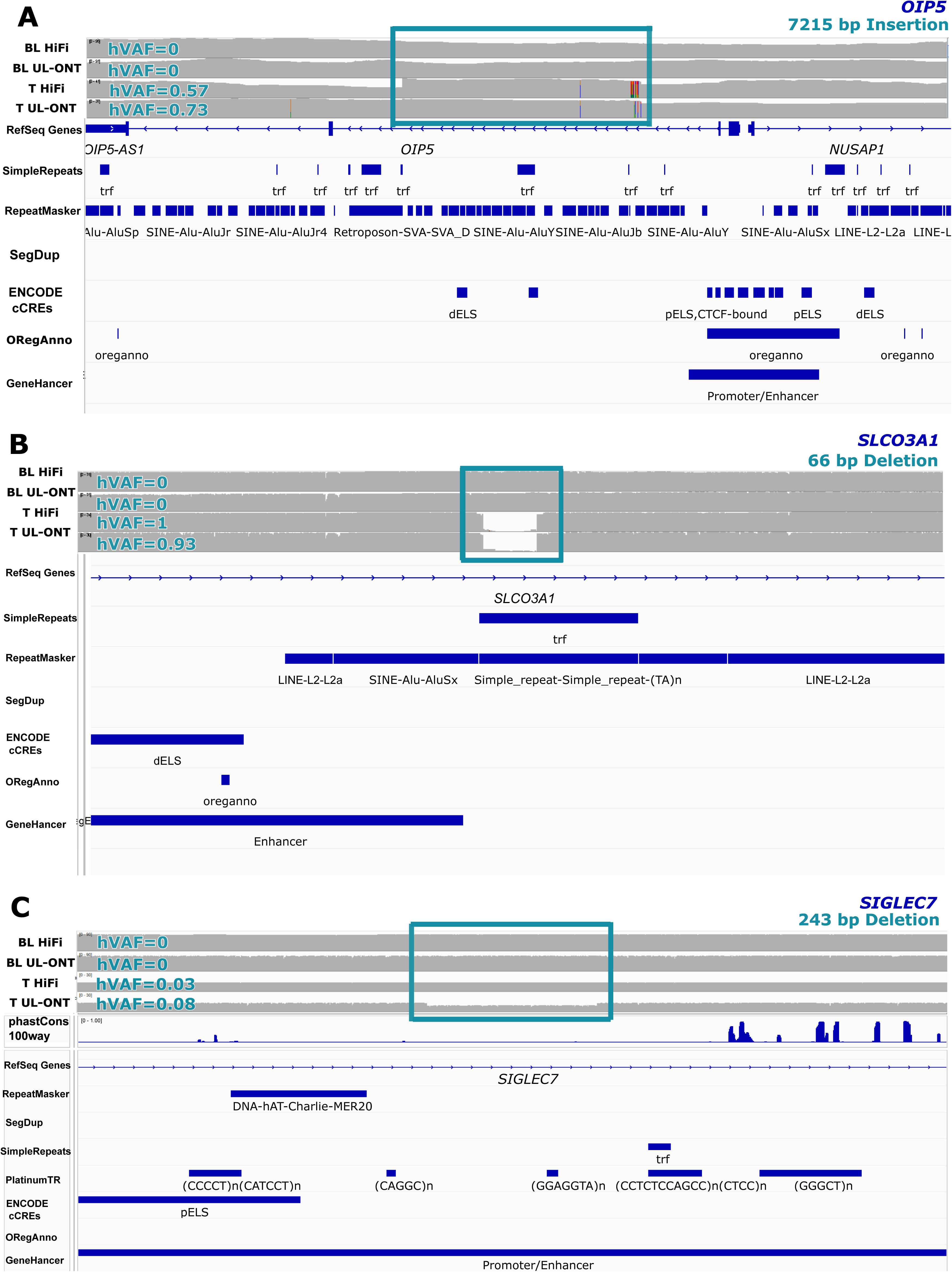
COLO829BL_DSA-only sSVs near regulatory regions. Examples of COLO829BL_DSA-only visually validated variants on one haplotype impacting **A)** *OIP5* (7,215 bp insertion), **B)** *SLCO3A1* (66 bp deletion), and **C)** *SIGLEC7* (243 bp deletion). Manually genotyped hVAF is shown for each platform. BL, blood; T, tumor.

## DISCUSSION

Here, we evaluated the utility of DSAs for detecting sSVs in the well-characterized COLO829 melanoma cell line, comparing results across standard reference genomes (GRCh38, CHM13) and both HiFi and ONT long-read sequencing platforms. We demonstrate that using COLO829BL_DSA captures approximately 1.8-fold more visually validated sSVs than either GRCh38 or CHM13. Importantly, this improvement reflects increased sensitivity both within genomic regions shared across references and within COLO829BL_DSA-specific sequence that is absent from standard reference assemblies. All COLO829BL_DSA-unique sSVs localize to a tandem repeat or segmental duplication region of the genome, which are notoriously difficult to capture with traditional references. Beyond the uniquely represented sequence, COLO829BL_DSA also improved sSV detection within regions shared across the references, identifying eight additional validated variants that were missed when using GRCh38 or CHM13. Together, these results highlight the value of individualized genome assemblies for more completely capturing the full landscape of somatic variation, particularly in complex and repetitive genomic regions.

We also provide practical suggestions for how to handle the complexities of working with DSAs. Because DSAs are diploid, reads originating from regions that are nearly identical between haplotypes can map ambiguously to both copies, leading to reduced mapping quality scores and dropout of true variants. Manually adjusting mapping quality scores for reads aligned to COLO829BL_DSA mitigates this, extending an approach previously demonstrated for somatic single-nucleotide variants (Sohn et al. 2026) to sSVs. This recovered most validated true positives that would otherwise have been missed, at a cost in precision concentrated in variants supported by fewer than all three callers. An alternative strategy analyzes each haplotype independently and merges calls downstream (Zhang et al. 2025). However, forcing each read to map to only one haplotype can cause mis-mapping, with artifacts concentrated in the most structurally divergent regions between haplotypes, exactly where accurate sSV detection matters most. Our approach avoids this by calling sSVs directly from the phased diploid assembly. Still, future callers that natively handle diploid, haplotype-resolved references, for example by mapping directly to a donor-specific graph, would likely reduce this tradeoff further.

Separately, we demonstrate the utility of pangenome graph-based methods to map sSVs from COLO829BL_DSA to common coordinate systems, such as CHM13. The high proportion of COLO829BL_DSA variants that could not be successfully lifted over, especially in satellite and repeat-rich regions, reinforces the idea that linear references often miss biologically relevant variation. This limitation extends to the pangenome graph itself. Reads supporting the 36 COLO829BL_DSA-unique variants failed to map, or mapped with very low confidence (MAPQ <5), to the graph, consistent with these loci falling in satellite sequence largely excluded from current graph construction due to the spurious edges such repetitive sequence introduces. While pangenome graphs are a valuable tool for coordinate conversion and cross-reference comparison, they do not substitute for a DSA in recovering this most repetitive fraction of the genome.

A central challenge of this work was balancing sensitivity against the scalability of validation. While 71% of the total consensus sSVs across reference genomes turned out to be false positives upon inspection, validation rates varied sharply by genomic context: 44–56% outside TR regions across the three references, but only 1–16% within them, where the DSA’s advantage is greatest. To move beyond visual review as the primary safeguard, we systematically compared caller, technology, and evidence-threshold combinations against our validated callset. We found that a region-specific strategy, requiring DELLY support from HiFi data outside TRs and Severus support from HiFi data within them, substantially improved precision over our original consensus criteria, and we recommend this approach for future consensus building. For studies without access to this specific caller/technology combination, requiring a minimum multi-caller evidence threshold (≥4) offers a more general second-best strategy. The higher error rate of UL-ONT likely explains its large contribution to raw and consensus candidate sSVs alongside its lower validation rate, motivating our reliance on cross-caller and cross-platform agreement rather than any single platform’s raw calls. For groups with cost constraints, HiFi alone offers a reasonable balance of sensitivity and precision, while multi-platform sequencing remains valuable for maximizing sensitivity, particularly in TR regions.

Our analysis also underscores the importance of using multiple sSV callers, while clarifying which combinations are worth the added complexity. No single tool comprehensively detected all high-confidence sSVs, and cross-caller support helped prioritize true somatic events, particularly for the low-VAF variants that are difficult to discriminate from sequencing noise. Among calls detected by only one caller across all sequencing data types, 69% of DELLY-only calls and 29% of Severus-only calls validated, while no Sniffles2-only calls did. Sniffles2-only calls can therefore be eliminated from future consensus datasets to reduce the burden of visual inspection without meaningful loss of sensitivity, and any Sniffles2-supported call should not be trusted without additional multi-caller corroboration.

Severus’s advantage in TR regions is likely due to specific preprocessing steps in its algorithm. Severus harmonizes inconsistently placed indels within variable-number tandem repeats before clustering (Keskus et al. 2026). DELLY and Sniffles2 do not apply comparable harmonization, which likely explains their weaker performance in these same regions. Extending this normalization to the larger, higher-order repeats characteristic of satellite sequence is a promising direction for future caller development.

We further observed that >95% of variants in the consensus dataset detected exclusively in either HiFi or ONT sequencing data, but not supported across both technologies, failed visual validation, indicating that cross-platform support is a strong indicator of call reliability. However, each long-read technology has distinct strengths and limitations (Wang et al. 2025; Harvey et al. 2023), and strict enforcement of cross-platform support would come at the cost of sensitivity. Notably, applying such a criterion would have resulted in the missed cancer-associated gene fusion between *TMX3* and *CCDC102B* since this was only detected in UL-ONT data. This underscores the importance of balancing specificity and sensitivity and highlights the continued need for orthogonal validation and manual review, particularly for variants of potential functional relevance.

Our analysis identified 17 sSVs that map to known cancer-associated genes; 15 of these were detected by all three references, and two were only identified by COLO829BL_DSA: *OIP5* and *SIGLEC7*. *OIP5*, which is located at the centromere, resides in a genomic region that is particularly prone to being overlooked by sSV calling approaches based on traditional references. Functionally, OIP5 plays a key role in mitosis and is frequently overexpressed in tumor tissues (Pan et al. 2023). *SIGLEC7* encodes an inhibitory immune checkpoint receptor and can promote immune evasion in cancer (Benthami et al. 2025; Thangaraj et al. 2025). Notably, *SIGLEC7-*associated sSVs have not been reported in prior COLO829 analyses.

A limitation of this analysis is its emphasis on specificity over sensitivity. Our consensus callset recovered 85% of the 68 experimentally validated sSVs reported by Espejo Valle-Inclan et al. 2022 within genomic regions accessible to their reference-based approach. The 36 sSVs unique to COLO829BL_DSA, by contrast, fall entirely within satellite sequence absent from GRCh38 and CHM13 and would not have been detectable by their method regardless of sequencing depth, underscoring that the benefit of the DSA lies in extending detection into regions structurally inaccessible to non-haplotype-resolved approaches. Our validation strategy differs from that of Espejo Valle-Inclan et al., who combined five sequencing platforms with PCR- and Bionano-based orthogonal confirmation; we instead relied on visual inspection of aligned reads and cross-caller support across three long-read SV callers. Overall, while our approach may miss some sSVs, the variants it identifies are highly reliable, especially when integrating the improved consensus set criteria demonstrated in Figure 3.

In summary, we provide practical, actionable guidance for future studies. Where cost is a constraint, we recommend prioritizing HiFi sequencing. We also recommend applying region-specific caller and technology filtering rather than a uniform consensus threshold, excluding Sniffles2-only calls, adjusting mapping quality to recover sSVs from the DSA, and using pangenome graphs to convert coordinates between assemblies. These findings underscore the value of phased, haplotype-resolved assemblies for comprehensive sSV discovery and suggest that continued refinement of repeat-aware calling algorithms, alongside broader adoption of DSAs, could further improve detection of functionally relevant structural variation in cancer and other complex genomic contexts. Moving forward, efforts within the SMaHT Network will extend these analyses across multiple tissues and donors, providing a more complete understanding of the normal sSV landscape in humans.

## METHODS

### Sequencing Data Generation

Sequencing data for both COLO829 and its matched normal, COLO829BL, were sourced from the SMaHT benchmarking effort (Somatic Mosaicism across Human Tissues Network 2026) (RRID:CVCL_1137). Sequencing data were generated using PacBio HiFi, STD-ONT, and UL- ONT platforms (**Supplemental Table S1**). Comprehensive details for COLO829BL/COLO829 cell culture, library preparation, and sequencing are described in the SMaHT flagship paper (Coorens et al. 2025).The corresponding data can be accessed at https://data.smaht.org.

### Constructing the COLO829BL and COLO829 DSAs

To construct the DSA for COLO829BL, we used 60× coverage of PacBio HiFi and ONT data, as well as 30× Hi-C coverage. ONT reads were downsampled to enrich for the longest reads, and phased assembly was performed using Verkko (version 2.1) (Antipov et al. 2025). Post- assembly, we applied the same contamination filtering pipeline involving NCBI FCS (version 0.4.0 beta) (Astashyn et al. 2024), adapter screening, and BLAST-based identification and removal of mitochondrial, Epstein–Barr virus, and rDNA sequences. This process produced decontaminated, haplotype-resolved assemblies. Detailed methods for DSA construction and quality evaluation are described here (Sohn et al. 2026).

For the tumor assembly (COLO829), all available ONT sequencing data, including R9 and R10 chemistries as well as STD-ONT and UL-ONT reads, were used. The initial dataset had a read N50 of 60.041 kbp, with 24.788× coverage from reads longer than 100 kbp and a total coverage of 88.827×. To improve base-level accuracy, reads were processed using fastplong (version 0.2.2) (Chen 2023), applying both 5′ and 3′ trimming based on a mean read base quality threshold of Q20. After trimming, the dataset had a read N50 of 58.440 kbp, with 18.951× coverage from reads longer than 100 kbp and a total coverage of 66.632×.

A pseudo-phased assembly was generated using hifiasm (version 0.25.0) (--dual-scaf --telo-m CCCTAA) (Cheng et al. 2021). To remove potential contaminant sequences, including nonhuman sequences, Epstein–Barr virus, and residual adapter sequences, the assemblies were screened using NCBI FCS (version 0.4.0 beta) (Astashyn et al. 2024) in combination with BLAST (version 2.15) (Camacho et al. 2009).

### Alignment

All three types of raw sequencing reads (HiFi, STD-ONT, and UL-ONT), for both COLO829BL and COLO829, were aligned to each reference genome (GRCh38, CHM13, and COLO829BL_DSA) using pbmm2 (version 1.13.1) ((https://github.com/PacificBiosciences/pbmm2) for HiFi reads (pbmm2 align –preset CCS) and minimap2 (version 2.28) (Li 2018) for STD-ONT and UL-ONT reads (minimap2 -ax map-ont). To assess sequencing data quality across platforms, we calculated read length, base quality, and alignment error rate directly from the aligned COLO829 sequencing data using SAMtools stats (version 1.22) (Li et al. 2009). For each combination of sequencing platform (HiFi, STD-ONT, and UL-ONT) and reference genome (GRCh38, CHM13, and COLO829BL_DSA), we report the number of reads mapped, mean read length, mean base quality (Phred score), and error rate (**Supplemental Table S1**).

### Mapping Quality Score Adjustment

Aligning the sequencing reads onto the diploid COLO829BL_DSA would result in a mapping quality (MAPQ) score of zero, since the read can map to two different locations, representing each haplotype. This can result in variants being discarded in downstream sSV calling due to MAPQ filters by variant callers. To avoid this issue, we manually changed the MAPQ values for the COLO829BL_DSA reads to 60, as further described here (Sohn et al. 2026).

### sSV Calling

For each combination of COLO829BL or COLO829 data, type of sequencing data (HiFi, STD-ONT, and UL-ONT), and reference genome (GRCh38, CHM13, and COLO829BL_DSA), three different sSV callers were used to call sSVs. DELLY (version 1.5.0) (Rausch et al. 2012) somatic mode was used with the blood and tumor datasets run together and maximum VAF threshold set at 1. Severus (version 1.5) (Keskus et al. 2026) was used in tumor-normal mode with the blood and tumor datasets with the maximum VAF threshold set at 1. For Sniffles2 (version 2.7.1) (Smolka et al. 2024), COLO829BL and COLO829 data were run separately using both germline and mosaic mode and the resulting callsets were combined. Any variants detected in COLO829BL were removed from the COLO829 dataset using Truvari bench (70% sequence similarity and 500 bp distance) (English et al. 2022).

DELLY, Severus, and Sniffles2 were selected for this analysis based on their recall in a prior benchmarking study of long-read SV callers in cancer genomes (Aydin et al. 2025), with additional consideration given to tools planned for use by the SMaHT SV working group in the analysis of production donors, including Sniffles2.

### sSV Filtering

To remove artifacts from our raw sSV calls, we excluded variants called in regions of known artifacts, misassembly, or collapse for all three reference genomes. We exclude GRCh38 sSVs in gap or telomere regions (within 10 kbp to telomere) from UCSC Genome Browser (Perez et al. 2025), accounting for 218 Mbp. CHM13 sSVs in acrocentric short arms, centromeres, and satellite regions were excluded except for monomeric alpha-Sat (https://s3-us-west-2.amazonaws.com/human-pangenomics/T2T/CHM13/assemblies/annotation/chm13v2.0_censat_v2.1.bed) (Logsdon et al. 2025; Sui et al. 2026). We created a nonredundant BED file with BEDTools (Quinlan and Hall 2010), resulting in 246 Mbp of masked reference loci. These regions showed an excessive number of variants, specifically in human satellite repeat (e.g., HSat3, etc.), that are likely to be mapping artifacts. For COLO829BL_DSA, we removed any SVs called in regions of known misassembly or collapse. These regions were determined by NucFreq (https://github.com/mrvollger/NucFreq) and Flagger (Liao et al. 2023) (https://github.com/mobinasri/flagger). Then, to remove inherent germline differences between COLO829BL and GRCh38/CHM13, we ran PAV (Ebert et al. 2021) between the assemblies and removed any calls that overlapped from our downstream analyses.

### sSV Callset Integration

After the raw sSV callsets were created, variants were collapsed into one dataset per genome reference and caller regardless of sequencing data type. This process was done using Truvari default collapse parameters (70% sequence similarity, 500 bp distance, type matching not required) (English et al. 2022) on a merged VCF of the calls for HiFi, STD-ONT, and UL-ONT. Each of these collapsed datasets was then collapsed a second time with the same parameters to create one callset per reference genome using the same parameters, regardless of sSV caller. sSVs present in at least two out of three sSV caller datasets or three out of three sequencing data types for a single caller were considered high-confidence and are present in the consensus dataset.

### Visual Validation

SVs in the consensus set were visually evaluated through inspection of sequencing data and genome assemblies. Nucleotide frequency (NucFreq) (Vollger et al. 2019) plots were examined to assess local assembly quality and to identify regions with atypical base composition patterns that could indicate assembly errors. sSVs were further validated by visualizing aligned sequencing reads in the Integrative Genomics Viewer (IGV) (Robinson et al. 2011). sSVs were considered supported when read-level evidence in IGV was concordant with the predicted sSV and no obvious assembly artifacts were indicated by NucFreq analysis.

### Coordinate Conversion Between Assemblies

To compare the sSV calls between reference genomes, coordinates needed to be converted to a common coordinate system. Here we used CHM13. To convert the COLO829BL_DSA coordinates, we used a donor-specific pangenome graph constructed with Minigraph-Cactus (Hickey et al. 2024). It contains GRCh38, CHM13, and the two haplotypes of COLO829BL, allowing for bidirectional coordinate translation between assemblies and identification of COLO829BL_DSA-specific intervals. This process is described in more detail here (Sohn et al. 2026). For COLO829BL_DSA-unique variants that could not be converted to CHM13 or GRCh38 coordinates using this approach, we used SafFire (https://github.com/vollgerlab/SafFire) to determine their approximate syntenic position relative to these references, allowing their genomic context to be visualized despite the absence of a true corresponding coordinate.

### Identifying Tandem Repeat and Segmental Duplication Regions

Repeat element annotations were added using Rhondite (10.5281/zenodo.6036498), which combines RepeatMasker (version 4.1.5) (https://github.com/Dfam-consortium/RepeatMasker), Tandem Repeats Finder (TRF) (version 4.1.0) (Benson 1999), and DupMasker (Jiang et al. 2008) in a Snakemake (Mölder et al. 2021) workflow. Detailed methods are available here (Sohn et al. 2026).

### Segmental Duplication Annotation

Repetitive sequences in each genome assembly were identified and masked using three complementary tools. RepeatMasker (version 4.1.5) (https://github.com/Dfam-consortium/RepeatMasker) was run with the parameters “*-s -e ncbi -xsmall -species human [asm.fa]*” followed by TRF (version 4.1.0) (“*trf [asm.fa] 2 7 7 80 10 50 2000 -l 30 -h -ngs*”) (Benson 1999) and WindowMasker (“*windowmasker -mk_counts -mem 16384 -smem 2048 - infmt FASTA -sformat obinary -in [asm.fa] -out [asm.count]* && windowmasker -infmt FASTA - ustat [asm.count] -dust T -outfmt interval -in [asm.fa] -out [asm.interval]”) (Morgulis et al. 2006) to comprehensively mask the repeats. The repeat annotations from all three tools were integrated, and the merged BED intervals were used to softmask the assemblies. Segmental duplications were then identified on these masked assemblies using SEDEF (version 1.1) (Numanagić et al. 2018). For subsequent analyses, only duplications exceeding 1 kbp in length, with >90% sequence identity and <70% satellite DNA content, were retained.

### Identifying Gene Overlap and Annotations

To determine whether the visually validated sSVs overlapped annotated genes, sSV coordinates were compared against gene annotations using the UCSC Genome Browser (Perez et al. 2025) (https://genome.ucsc.edu/). sSV intervals were visualized and queried within CHM13 and/or GRCh38 and overlap with known genes was assessed based on UCSC gene annotation tracks. An sSV was considered gene-overlapping if any portion of the variant interval intersected the genomic coordinates of an annotated gene, including exonic or intronic regions. To identify SV location in relation to exons/introns, promoters/enhancers, and regulatory elements, we integrated published datasets from UCSC Genome Browser tracks, including candidate cis-regulatory elements (ENCODE Regulation and ENCODE cCREs [ENCODE Project Consortium et al. 2020], ORegAnno [Lesurf et al. 2016], GeneHancer [Fishilevich et al. 2017]) and ENCODE histone marks (H3K27ac, H3K4me1, H3K4me3; ENCODE Project Consortium et al. 2020).

## DATA ACCESS

The SMaHT COLO829 benchmarking data can be accessed from the SMaHT Data Portal at https://data.smaht.org/. SMaHT benchmark core-assay data are released on dbGaP (<u>phs004193</u>). Both are available for those outside of the Network to apply for access.

## ACKNOWLEDGEMENTS

This work was supported by the NIH Common Fund through the Office of Strategic Coordination/Office of the NIH Director under awards U24 MH133204, U24 NS132103, UG3 NS132024, UG3 NS132061, UG3 NS132084, UG3 NS132105, UG3 NS132127, UG3 NS132128, UG3 NS132132, UG3 NS132134, UG3 NS132135, UG3 NS132136, UG3 NS132138, UG3 NS132139, UG3 NS132144, UG3 NS132146, UM1 DA058219, UM1 DA058220, UM1 DA058229, UM1 DA058230, UM1 DA058235, and UM1 DA058236. A.B.S. is supported by a Career Award for Medical Scientists from the Burroughs Wellcome Fund and is a Pew Biomedical Scholar. Research reported in this publication was supported, in part, by the National Human Genome Research Institute of the National Institutes of Health (NIH) under Award Numbers 1DP5OD029630 and 1U01HG013744 to A.B.S. and R01HG010169 to E.E.E. The content is solely the responsibility of the authors and does not necessarily represent the official views of the NIH. NIH training grants supported M.R.V. and S.C.B. (T32GM007454) and T.M. (T32HG000035). M.R.V. additionally received support from a Pathway to Independence Award from the National Institute of General Medical Sciences (1K99GM155552–01). E.E.E. is an investigator of the Howard Hughes Medical Institute.

The study was conceived by T.M, L.R., M-H.S., A.B.S., & E.E.E. Statistical analyses were performed by T.M., J.L., L.R., M.S., A.M., Y.K., D.Y., Y.S., & F.K.M. Data generation, processing, quality control, and organization were performed by K.M.M., K.H., M.S., M.A., K.A.S., N.K., J.O., M.D.N., A.S-C., A.L., C.N.J., C.O., C.D.F., C.Z., D.M.J., E.G.S., E.R., J.T.K., J.R., L.S., M.R.V., K.L., M.M.P., M-F.H., N.YT.A., P.M.N., S.R.M., S.N., S.B., T.S., V.F., Y.M., B.C.S., M.R., J.D.S., & J.M.W. The manuscript was written by T.M. with substantial input from E.E.E., J.L., M-H.S., Y.K., D.Y., & Y.S. and critical revisions from all authors. Senior supervision was provided by N.L.P., C-L.W., J.T.B., A.B.S, & E.E.E. All authors reviewed and approved the final manuscript.

