## Supplementary Material: Figs S1-S2 & Tables S1-S2 for "Donor-specific assemblies enhance somatic structural variant detection in complex genomic regions"

### Table of contents

Supplemental Fig. S1. Raw sSV calls and filtering p. 2

Supplemental Fig. S2. Platform and caller evidence for visually validated sSVs p. 3

Supplemental Table S1. Sequencing quality metrics for COLO829 data p. 4

Supplemental Table S2. Visually validated sSVs in sequence unique to the DSA p. 5

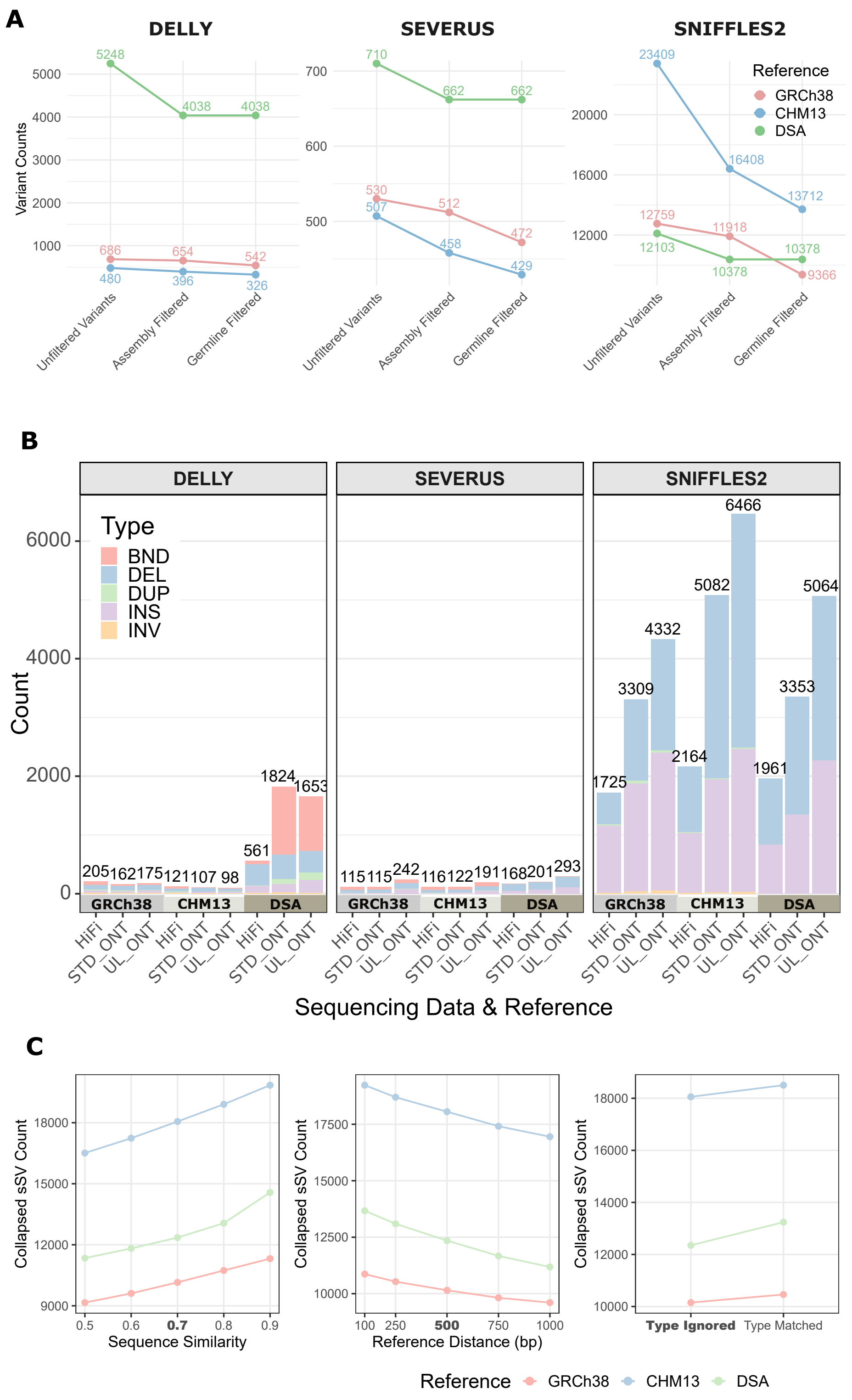


**Supplemental Fig. S1. Raw sSV calls and filtering. A)** Post-sSV calling filtering to remove germline and sequencing artifacts, including calls overlapping misassembled, collapsed, or known artifact regions across all three references ("Assembly Filtered"), with additional PAV-based removal of germline artifacts arising from inherent DSA versus GRCh38/CHM13 reference differences ("Germline Filtered"), retaining variants used for downstream analyses. **B)** The calls that remained after the initial filtering stratified by sSV type, reference genome, and sSV caller. In this depiction, the y-axis is constant across sSV callers. **C)** Sensitivity of the initial Truvari collapse step to variation in sequence similarity threshold (left), reference distance threshold (center), and SV type-matching requirement (right), shown for each reference genome. Value used for the study is bold.

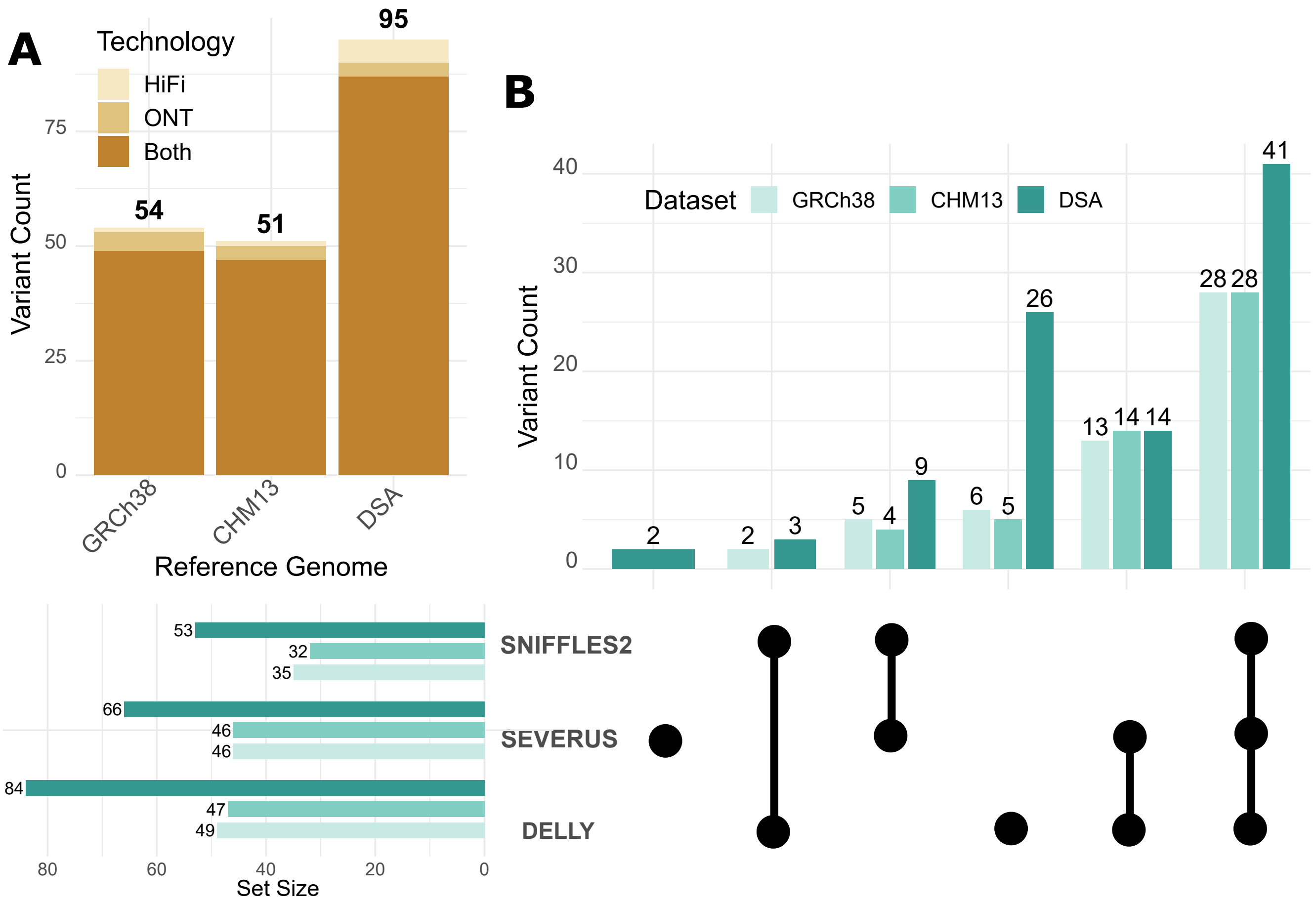


**Supplemental Fig. S2. Platform and caller evidence for visually validated sSVs. A)** Sequencing data types that called each true positive variant across HiFi and ONT. **B)** Combinations of sSV callers that called each true positive variant.

**Supplemental Table S1.** Sequencing quality metrics for COLO829 data.

| Reference | Platform | Reads Mapped | Mean Read Length (bp) | Mean Base Quality (Phred) | Error Rate |
|---|---|---|---|---|---|
| GRCh38 | HiFi | 8,916,884 | 21,579 | 37.4 | 1.2% |
| GRCh38 | STD ONT | 10,328,013 | 12,831 | 31.5 | 4.6% |
| GRCh38 | UL ONT | 3,042,422 | 41,201 | 27.7 | 5.3% |
| CHM13 | HiFi | 8,992,223 | 21,571 | 37.4 | 0.8% |
| CHM13 | STD ONT | 10,310,156 | 12,831 | 31.5 | 3.8% |
| CHM13 | UL ONT | 3,042,399 | 41,201 | 27.7 | 4.5% |
| COLO829BL_DSA | HiFi | 8,992,037 | 21,571 | 37.4 | 0.3% |
| COLO829BL_DSA | STD ONT | 10,329,457 | 12,831 | 31.5 | 3.0% |
| COLO829BL_DSA | UL ONT | 3,046,730 | 41,201 | 27.7 | 3.7% |

**Supplemental Table S2.** Visually validated sSVs in sequence unique to the DSA.

| Contig | Start | End | Type | Length | Genotype |
|---|---|---|---|---|---|
| haplotype1-0000001 | 93927875 | 93930035 | DEL | 160 | 1/1 |
| haplotype1-0000002 | 96236768 | 96240634 | INS | 1866 | 0/1 |
| haplotype1-0000002 | 97440643 | 97443323 | DEL | 680 | 0/1 |
| haplotype1-0000003 | 101293834 | 101297028 | DEL | 1194 | 0/1 |
| haplotype1-0000005 | 6623250 | 6627978 | DEL | 2728 | 0/1 |
| haplotype1-0000006 | 34914884 | 34921985 | DEL | 5101 | 0/1 |
| haplotype1-0000009 | 97535520 | 97538542 | DEL | 1022 | 0/1 |
| haplotype1-0000012 | 124992387 | 124996007 | DEL | 1620 | 0/1 |
| haplotype1-0000012 | 125630546 | 125632886 | INS | 340 | 1/1 |
| haplotype1-0000012 | 131676934 | 131680710 | DEL | 1776 | 0/1 |
| haplotype1-0000012 | 132331383 | 132334017 | DEL | 634 | 0/1 |
| haplotype1-0000012 | 138299844 | 138302388 | DEL | 544 | 0/1 |
| haplotype1-0000012 | 138401758 | 138404201 | DEL | 443 | 1/1 |
| haplotype1-0000012 | 140488090 | 140493174 | DEL | 3084 | 0/1 |
| haplotype1-0000012 | 142654253 | 142656721 | DEL | 468 | 0/1 |
| haplotype1-0000015 | 61667959 | 61670301 | DEL | 342 | 0/1 |
| haplotype1-0000017 | 52337179 | 52340029 | DEL | 850 | 0/1 |
| haplotype1-0000018 | 7882286 | 7885650 | DEL | 1364 | 0/1 |
| haplotype1-0000019 | 94142349 | 94146390 | INS | 2041 | 0/1 |
| haplotype1-0000021 | 35349405 | 35352085 | DEL | 680 | 0/1 |
| haplotype1-0000026 | 60996007 | 60998657 | DEL | 650 | 0/1 |
| haplotype1-0000030 | 161997 | 171711 | DEL | 7714 | 0/1 |
| haplotype2-0000050 | 9064342 | 9069355 | DEL | 3013 | 1/1 |
| haplotype2-0000050 | 10567817 | 10570280 | INS | 463 | 1/1 |
| haplotype2-0000051 | 99918999 | 99924060 | DEL | 3061 | 0/1 |
| haplotype2-0000055 | 57433013 | 57437727 | DEL | 2714 | 0/1 |
| haplotype2-0000056 | 98038927 | 98041949 | DEL | 1022 | 0/1 |
| haplotype2-0000056 | 98639926 | 98642947 | DEL | 1021 | 0/1 |
| haplotype2-0000058 | 107362225 | 107364519 | DEL | 294 | 0/1 |
| haplotype2-0000061 | 47130985 | 47134857 | DEL | 1872 | 0/1 |
| haplotype2-0000065 | 8723572 | 8726937 | DEL | 1365 | 1/1 |
| haplotype2-0000071 | 2693586 | 2697130 | DEL | 1544 | 0/1 |
| haplotype2-0000071 | 2934242 | 2936988 | DEL | 746 | 0/1 |
| haplotype2-0000072 | 35311540 | 35315575 | INS | 2035 | 0/1 |
| haplotype2-0000072 | 36405620 | 36408975 | DEL | 1355 | 0/1 |
| haplotype2-0000072 | 36421268 | 36425646 | DEL | 2378 | 0/1 |
